# Cardiac Localized Polycystin-2 plays a Functional Role in Natriuretic Peptide Production and its Absence Contributes to Hypertension

**DOI:** 10.1101/2024.01.02.573922

**Authors:** Brandon Elliott, Karla M. Márquez-Nogueras, Paula Thuo, Elisabeth DiNello, Ryne M. Knutila, Geena E. Fritzmann, Monte Willis, Arlene B. Chapman, Quan Cao, David Y. Barefield, Ivana Y. Kuo

**Author notes:** **Address for correspondence**: Ivana Y. Kuo.

## Abstract

Cardiovascular complications are the most common cause of mortality in patients with autosomal dominant polycystic kidney disease (ADPKD). Hypertension is seen in 70% of patients by the age of 30 prior to decline in kidney function. The natriuretic peptides (NPs), atrial natriuretic peptide (ANP) and brain natriuretic peptide (BNP), are released by cardiomyocytes in response to membrane stretch, increasing urinary excretion of sodium and water. Mice heterozygous for *Pkd2* have attenuated NP responses and we hypothesized that cardiomyocyte-localized polycystin proteins contribute to production of NPs. Cardiomyocyte-specific knock-out models of polycystin-2 (PC2), one of the causative genes of ADPKD, demonstrate diurnal hypertension. These mice have decreased ANP and BNP expression in the left ventricle. Analysis of the pathways involved in production, maturation, and activity of NPs identified decreased transcription of CgB, PCSK6, and NFAT genes in cPC2-KOs. Engineered heart tissue with human iPSCs driven into cardiomyocytes with CRISPR/Cas9 KO of *PKD2* failed to produce ANP. These results suggest that PC2 in cardiomyocytes are involved in NP production and lack of cardiac PC2 predisposes to a hypertensive volume expanded phenotype, which may contribute to the development of hypertension in ADPKD.

## INTRODUCTION

Autosomal dominant polycystic kidney disease (ADPKD) is the most common hereditary kidney disease, occurring in 1 in 700 births [1]. Patients develop bilateral kidney cysts, leading to progressive loss of kidney function ultimately requiring renal replacement therapy [1,2]. Cardiovascular complications are the most common cause of mortality of ADPKD patients [3–7] and hypertension is one of the earliest detectable symptoms of ADPKD, occurring in approximately ∼70% of ADPKD patients before the age of 30. Long standing hypertension is a significant contributor to cardiovascular disease in ADPKD [8–12]. Given the progressive nature of ADPKD, slowing decline in kidney function, and controlling cardiovascular risk factors such as hypertension, are critical.

Increased activation of the renin-angiotensin-aldosterone system (RAAS) and endothelial nitric oxide dysregulation have been demonstrated to be major contributors to hypertension in ADPKD [13–15]. However, it is unclear if there is a primary cardiac defect due to mutations in the PKD genes that contribute to elevation in blood pressure and the development of hypertension. Importantly, elevations of RAAS is present in ADPKD individuals, but overall, the RAAS is relatively suppressed until sodium restriction occurs which then leads to an elevation in RAAS activation, very similar to bilateral artery stenosis where reductions in sodium and water excretion are seen. It is possible that defects in natriuresis exist in ADPKD and the examination of polycystin proteins in cardiovascular functioning may provide possible insight into this matter.

Mutations in *PKD1* and *PKD2*, which encode the proteins polycystin 1 (PC1) and polycystin 2 (PC2), account for ∼82% and ∼13% of ADPKD cases respectively, with other genetic mutations being rare [1]. PC1 and PC2 are transmembrane proteins that interact [16] to mediate downstream signaling processes, with PC2 forming a non-selective cation channel [17]. PC1 and PC2 reside in the endoplasmic reticulum, primary cilium, and the plasma membrane [18–21]. Loss of the signaling pathways regulated by the polycystin proteins result in kidney cysts. Polycystin proteins are also found in cardiomyocytes, with PC1 associated with potassium channels and the L-type Ca^2+^ channel [22,23] while PC2 associates with the ryanodine receptor [24,25]. The polycystins, either singly or in complex are mechanosensors in the kidney and cardiac settings [22,26–29], and the heterozygous PC2 mouse has impaired activation of the natriuretic peptide (NP) pathway under conditions that promote increased intravascular pressure and volume overload [30].

Decreases in extracellular osmolality or intravascular volume increases results in the cardiac production of NPs. Atrial natriuretic peptide (ANP) and brain natriuretic peptide (BNP) are produced primarily in the cardiac atria and ventricles respectively [31,32] in the setting of excessive cardiomyocyte stretch [33,34]. Cardiac NPs act on natriuretic peptide receptors (NPRs) in the nephron, increasing excretion of sodium with water [35]. Previous literature has shown that mice heterozygous for PC2 have an impaired NP production, in response to beta adrenergic stress or pressure overload [30]. Additionally, NPs increase glomerular filtration rate (GFR) by dilation of the afferent arteriole and constriction of the efferent arteriole [36] and also directly inhibit renin release from the kidney [37]. NP dysfunction impairs natriuresis leading to an increased circulating blood volume, elevations in blood pressure and hypertension, as evidenced by knockout of NP molecules or their regulators [38–40]. Despite the significance of the NP signaling pathways, the precise mechanistic signaling pathway from mechanical stretch to NP production currently is unclear.

As activation of the NP pathway is associated with a rise in intracellular Ca^2+^ [41–45], and PC2 has been demonstrated to both flux and modulate Ca^2+^ signaling, we hypothesized that lack of cardiomyocyte-localized PC2 could contribute to hypertension due to a lack of NP production. To test our hypothesis, we used an inducible cardiomyocyte specific knockout of PC2 (cPC2-KO) to isolate the role of PC2 in cardiomyocyte function. cPC2-KO mice had diurnal hypertension, impaired ANP and BNP mRNA production, and decreased transcription of genes needed for functional maturation and activity of the NPs. Human iPSC cells driven into cardiomyocytes with CRISPR/Cas9 KO of *PKD2* demonstrated diminished ANP production. Finally, a pathological PC2 mutant that cannot flux Ca^2+^ and full length PC2 both restored ANP production in PC2 deficient myoblast cells. These results suggest that PC2 acts specifically in cardiomyocytes to modulate the Ca^2+^ signal required for NP production.

## METHODS

### Animal Model

A cardiomyocyte inducible PC2 KO (cPC2-KO) mice were generated by crossing *Pkd2* floxed mice (gift from S. Somlo, Yale University) with αMHC-MerCreMer (Jackson Laboratories) mice (C57-Bl/6 background), as previously described [24]. Some mice were further crossed with gCamp6F-tdTomato mice (Salsa6f mice) [46]. Expression of this inducible cardiomyocyte ratiometric indicator allows for the measurement of cytosolic calcium. Starting at 8 weeks of age, mice were fed a tamoxifen chow diet (250mg·kg^-1^·g^-1^) for 10-14 days, resulting in Cre expression in cardiomyocytes and the respective cardiac specific knockout of *Pkd2.* Both male and female mice were used in subsequent experiments. All animal procedures and studies were performed under Institutional Animal Care and Use Committee (IACUC) protocols approved at Loyola University Chicago.

### Ultrasound of the kidney and heart

In a separate cohort of mice, ultrasound images (B-mode long-axis and M-mode short axis) were taken in mice 4 months after cPC2-KO (i.e.: 4 months of age, and again, 3 months after knockout, around 7 months of age) age on a VevoSonics 3100. Using VevoLab software, the EF was measured from 3 cardiac cycles from images taken in M-mode.

### Telemetry

At 20 weeks of age (i.e.: 4 months after cPC2-KO), catheters were implanted in mice via the carotid artery to measure blood pressure. Following 14 days of recovery, blood pressures are remotely recorded on a regular schedule (every 1-2 h for ∼2 min) using the TSE software. Data are analyzed for mean arterial pressure (MAP), heart rate (HR), and night, and day diastolic and systolic measurements were averaged for up to 2 weeks (to a final age of 7 months, or 5 months after cPC2-KO).

### Tissue harvest

At the completion of the study, mice were euthanized with isoflurane and retro orbital bleeds performed with heparinized capillary tubes to obtain serum for metabolic profiling. The mouse was then perfused with saline via the left ventricle, as previously described [24]. The heart was removed, and samples of the left ventricle was taken for protein studies (immediately frozen in liquid nitrogen), mRNA analysis (sample collected in RNAlater for RNA isolation) or immunofluorescence studies (places in 2% paraformaldehyde, PFA for 1 h).

### Protein studies

Protein was isolated from the left ventricle as previously described [24]. In brief, samples were homogenized in RIPA buffer containing protease and phosphatase inhibitors. 15-20µg of isolated protein was separated on SDS-reducing PAGE gels and transferred to PVDF membranes. The following antibodies were then used: p-AKT (1:1,000; Cell Signaling), AKT (1:1,000; Cell Signaling), p-GSK3ϕ3 (1:1,000; Cell signaling), GSK3ϕ3 (1:1,000; Cell signaling), p-CREB (Ser133; 1:1,000; Cell signaling), CREB (1:1,000; Cell signaling).

### mRNA studies

RNA was isolated from the left ventricle using the Direct-zol RNAminiprep kit (Zymo Research), following manufacturer’s instruction, and reverse transcribed into cDNA with random hexamer primers (Biobasics), following manufacturer’s instructions. Quantitative PCR reactions with SYBR Green technology with 20ng cDNA were conducted for the following gene targets: NPPA, NPPB, GATA4, Chromogranin B, NFκϕ3, PCSK6, Corin, NPR1, NPR2, NFATC2 and NFATC4. All reactions were conducted in duplicate, with GAPDH as housekeeping gene. All reactions were run on a QuantStudio platform (Fisher Scientific).

### Ex-vivo measurements of Ca^2+^

αMHC Mer-cre-mer, and αMHC Mer-cre-mer *Pkd2*^f/f^ mice were crossed with the Salsa6f mice to generate cardiomyocyte Salsa6f expressing mice after induction with tamoxifen [47]. The hearts were cannulated via the aorta and mounted on a modified Langendorff perfusion system. Hearts were perfused at 36°C (inline heater, Warner) with Tyrodes’ solutions (in mM: 140 NaCl, 4 KCl, 2 MgCl2, 5 EGTA, 10 glucose, 10 HEPES, pH 7.4) containing (S)-blebbestatin (25 μM; Cayman Chemicals) to minimize motion artifacts. The right atrium was imaged with a modified chamber using a 2.5x objective lens mounted on a Zeiss inverted Axioscope. The heart was alternatively excited with 488 and 543 LEDs and images acquired with a Flash 4 ORCA camera (Hamamatsu) at 20ms. Isoproterenol (10 nM) or Tyrodes’ solution adjusted to 200mOsm (by removing NaCl but keeping all other salt concentrations the same) were introduced via the perfusion system.

### Immunofluorescence microscopy

Following fixation of the left ventricle in 2% paraformaldehyde (1hr), samples were washed in PBS and impregnated with 30% sucrose followed by optimal cutting temperature (OCT) media. Sections (12-14µm) were cut on a cryostat (Leica) and air dried. Sections were blocked with 2% bovine serum albumin (BSA) and permeabilized with 0.2% triton-x. The following primary antibodies were then applied: GATA-4 (1:400; Cell Signaling), atrial natriuretic peptide (1:100; Santa Cruz), B-type natriuretic peptide (1:100; Santa Cruz), polycystin 2 (1:100; Santa Cruz). Primary antibodies were detected with fluorescent secondary antibodies (Alexa Fluor 488, Rhodamine and Cy-3). Sections were counterstained with phalloidin labelled with AF-647 (Cayman Chemicals) and mounted in pro-long diamond with DAPI (Thermo Fisher Scientific). Images were acquired on a Zeiss 880 microscope with an Airyscan detector with Zen software.

### Cell culture

C2C12 cells with either a no-template control or with PC2 knocked out by CRISPR/Cas9 technology were cultured in DMEM with 10% FBE and 5% penicillin/streptomycin at 37°C in a 5% CO_2_ humidified incubator following established procedures [48]. PC2-KO cells were transfected with constructs containing full-length mCherry labelled PC2 or mCherry labelled PC2 with a single point mutation (D511V) using PEI as a transfection agent. Cells were incubated in DMEM with 10% FBE or DMEM with 2% FBE for 3 days. RNA was isolated (Direct-zol RNAminiprep, Zymo Research) followed by reverse transcription into cDNA (Biobasics). Quantitative PCR experiments were then conducted as described above.

### IPSCs

Human iPSCs were differentiated into cardiomyocytes following previously established protocols [49]. Following differentiation, as evidenced by spontaneous beating, ∼1.5 million cells were seeded into engineered heart tissue constructs (Propria) [50–53]. Lentivirus containing guide sequences to either *PKD2* or a no-template control sequence combined with Cas9 was applied to the constructs at 250 MOI. Successful incorporation of the virus was evidenced by Zeogreen fluorescence. 3 days after virus transduction, constructs were mounted on the myopacer (Propria). Length dependent activation (LDA) protocols were conducted [50,54], starting with a –5% stretch up to a 5% stretch in Tyrode’s solution followed by Tyrode’s solution an titrated to an osmolarity of 270mOsm (by decreasing NaCl, but keeping all other osmotic components the same). Following the completion of the experiment, the myopod construct was fixed in PFA for immunofluorescent assays.

### Metabolomics

Plasma samples were assayed using the MxP 500 Quant kit (Biocrates) at the Indiana University Metabolomics Core.

### Statistical analysis

Data was plotted using GraphPad PRISM 9. We assumed non-parametric distributions where n was less than 15. Comparison of 2 independent groups were analyzed by Mann-Whitney U (non-parametric) and student’s t-test (parametric). Comparison of multiple groups were analyzed through Kruskal-Willis followed by Dunn’s multiple comparisons (non-parametric) or 1-way ANOVA followed by Sidak’s multiple comparisons (parametric) depending on the experimental approach. Conditions were considered statistically significant when p-values were <0.05. Error bars indicate Standard error of the mean (SEM). P-values are listed in each figure.

## RESULTS

### cPkd2-KO mice have elevated blood pressure

Blood pressure and cardiac function were measured four months after gene deletion (Fig. 1A). Mean arterial pressure (MAP), (systolic and diastolic blood pressures) were significantly elevated in all animals of cPC2-KO (Fig. 1B). When delineating between systolic and diastolic blood pressures (Fig. 1C), increases in both parameters were seen across both sexes in the cPC2-KO mice. There was no significant change in heart rate between animal groups (Fig. 1D).

**FIGURE 1:**
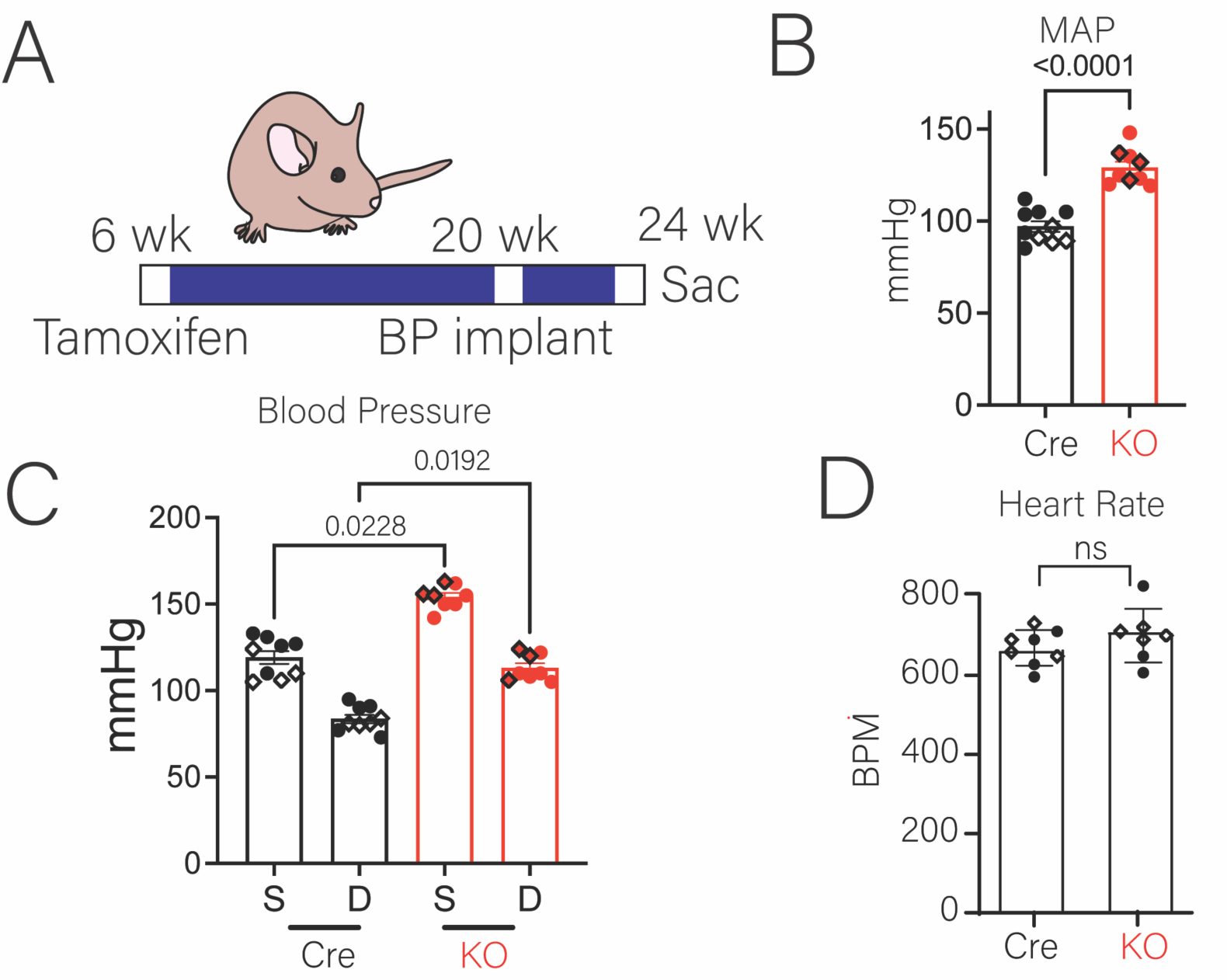
Cardiomyocyte specific PC2 Knockout Mice demonstrate elevated blood pressure without gross cardiac dysfunction. **A.** Diagram of experimental approach in both control and cPC2-KO mice. **B.** Mean arterial pressure (MAP) was significantly increased in both cPC2-KO female and male mice. Dots represent individual mice, with circles representing males, and diamonds representing females. Bars represent mean±SEM. Statistical analysis was done using Mann-Whitney U test. P-values are listed in the figure. **C.** Systolic (S) and Diastolic (D) blood pressure was significantly increased in both cPC2-KO female and male mice. Dots represent individual mice, with circles representing males, and diamonds representing females. Bars represent mean±SEM. Statistical analysis was done using Kruskal Willis followed by Dunn’s. P-values are listed in the figure. **D.** No change in heart rate between control and cPC2-KO mice in both female and male mice. Dots represent individual mice, with circles representing males, and diamonds representing females. Bars represent mean±SEM. Statistical analysis was done using Mann-Whitney U test. P-values are listed in the figure.

### Cardiac function is preserved in cPC2-KO mice

There were no differences in ejection fraction between the WT and cPC2-KO cohorts (Supplemental Fig. 1B). Left ventricular mass was not different between groups (Supplemental Fig. 1C). Global measures of radial and longitudinal strain and the inner diameter of the left ventricle in diastole were not different between the two groups and there were no sex differences (Supplemental Fig.1D, E, F). The ventricular posterior wall was increased in females and in males during systole (Supplemental Fig. 1G). Overall, there was no cardiac hypertrophy in the knockout mice. In these cardiac specific cPC2-KO mice there were no kidney cysts, increase in kidney volume or increases in renin expression (Supplemental Fig. 2A-G). We also examined by metabolomics the profile of over 300 substances in the plasma (Supplemental extra data set). We specifically examined the citrulline pathway due to its connection to nitric oxide and found that there was no significant difference in this pathway (Supplementary Fig. 3).

### Impaired production of Nppa and Nppb

To understand why cardiomyocyte specific loss of PC2 resulted in hypertension, we examined NP expression (Fig. 2A). As NPs function to decrease circulating blood volume, we expected to see upregulation of their transcription in the setting of elevated blood pressure in cPC2-KO mice. Instead, we saw impaired ANP and BNP gene transcription, where normalized expressions for *Nppa* (Fig. 2B) and *Nppb* (Fig. 2C) were significantly decreased in cPC2-KO mice compared to their WT counterparts. The AKT-GSK3β-GATA4 pathway which upregulates NPs in the setting of beta-adrenergic stress were unchanged. Left ventricular tissue protein expression of phosphorylated and total amounts of the kinase AKT, showed no difference between WT and cPC2-KO mice (Fig. 2D). Downstream of pAKT is GSK3β which relative to unphosphorylated GSK3β, did not differ between WT and cPC2-KO mice (Fig. 2E). The mRNA levels of the transcription factor GATA4 were also similar between WT and cPC2-KO mice (Fig. 2F). As the functional activity of GATA4 is dependent on its nuclear localization, we performed immunofluorescence assay of left ventricular tissue (Fig. 2G). We saw no difference in nuclear localized GATA4 between WT and cPC2-KO mice (Fig. 2H).

**FIGURE 2:**
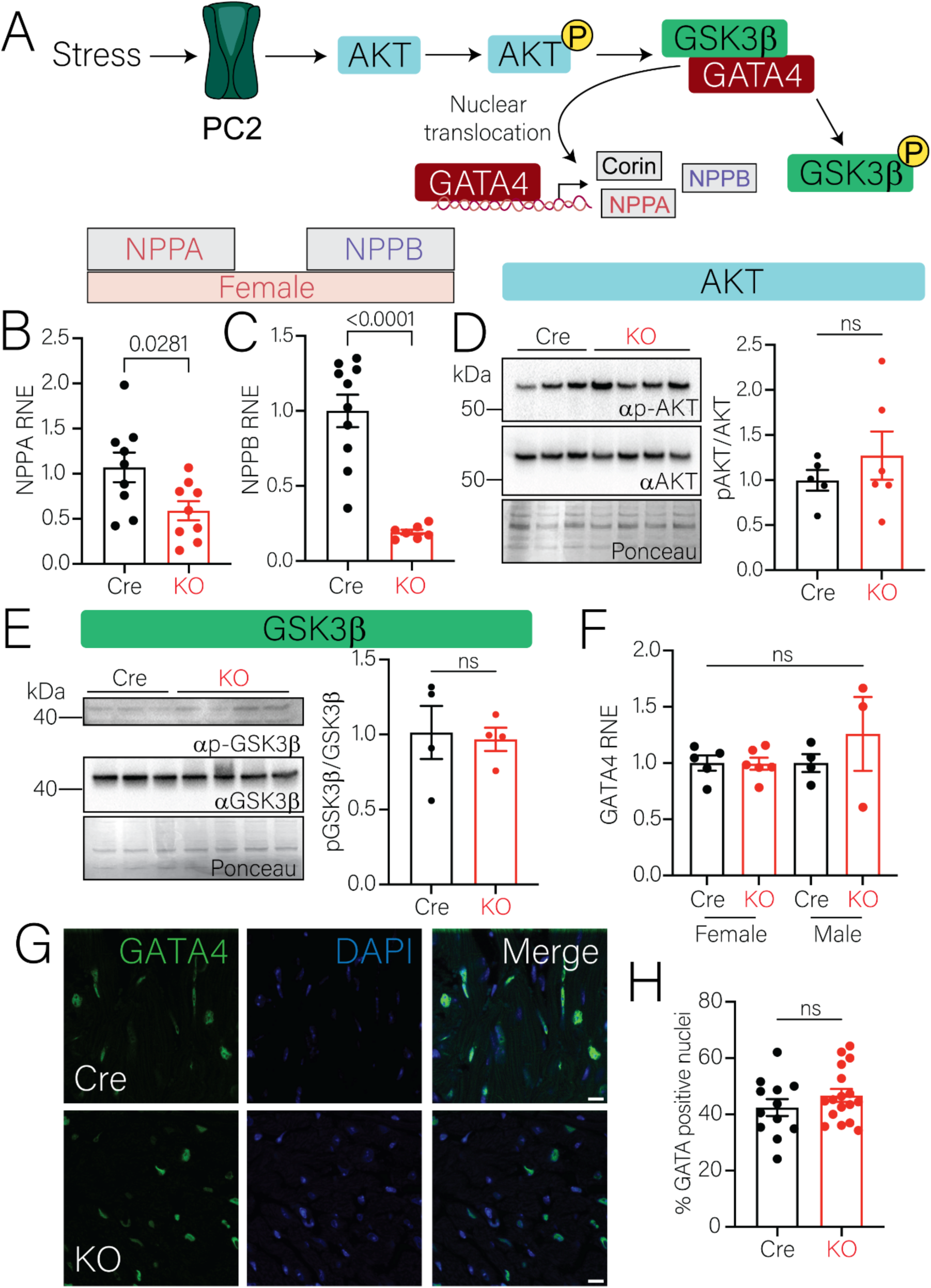
Production of natriuretic peptides is decreased in cardiomyocyte specific PC2-KO mice. **A.** Diagram of natriuretic peptide production. **B.** Normalized mRNA expression of *Nppa* is significantly decreased in left ventricle tissue from female cPC2-KO mice. Dots represent individual mice. Bars represent mean±SEM. Statistical analysis was done using Mann Whitney U test. p-values are listed in the figure. **C.** Normalized mRNA expression of *Nppb* is significantly decreased in left ventricle tissue from female cPC2-KO mice. Dots represent individual mice. Bars represent mean±SEM. Statistical analysis was done using Mann Whitney U test. p-values are listed in the figure. **D.** No significant change was observed in the expression of p-AKT/AKT ratio in cPC2-KO mice. Total protein was used as loading control. Dots represent individual mice. Bars represent mean±SEM. Statistical analysis was done using Mann Whitney U test. **E.** Western blot expression of p-GSK3β/GSK3β was unchanged between control and cPC2-KO mice. Total protein was used as loading control. Dots represent individual mice. Bars represent mean±SEM. Statistical analysis was done using Mann Whitney U test. **F.** mRNA expression of GSK3β was unchanged between control and cPC2-KO mice. Dots represent individual mice. Bars represent mean±SEM. Statistical analysis was done using student t-test. **G.** Representative immunofluorescent images from control and cPC2-KO mice with nuclear GATA4 staining. Scale bars represent 10μm. **H.** Quantification of GATA4 nuclei staining was unchanged between control and cPC2-KO mice. Dots represent individual cells analyzed from different tissues isolated from control and cPC2-KO mice. Bars represent mean±SEM. Statistical analysis was done using Mann Whitney U test.

### Reduction of processing enzymes, receptors, and downstream signaling for NP activity

BNP transcription is upregulated in response to beta adrenergic stress, possibly via a Chromogranin B (CgB) dependent mechanism, which is deficient in PC2 heterozygotes (Fig. 3A) [30]. Previous work has suggested that CgB is activated by Ca^2+^ [45,55]. Additionally, CgB is believed to be a constituent protein of exocytic vesicles containing ANP, and that its calcium sensing contributes to their release [56]. We found near complete abolishment of CgB mRNA expression in the left ventricles of cPC2-KO mice in both sexes, when compared to WT mice (Fig. 3B). We also looked at NFκβ, downstream of CgB in the BNP production pathway, and found a female-specific decrease in mRNA transcription from the left ventricle (Fig. 3C). Once mRNA of the NPs is produced, it must be processed. Pro-ANP is converted to ANP by the protease Corin [57], which itself is activated from pro-Corin to Corin via the protease PCSK6 [40] (Fig. 3D). Corin can also help to process BNP [58,59]. We found significantly decreased PCSK6 mRNA expression from left ventricular tissue of cPC2-KO mice compared to WT mice, in both sexes (Fig. 3E) while corin mRNA was not found to be different between WT and cPC2-KO mice (Fig. 3F).

**FIGURE 3:**
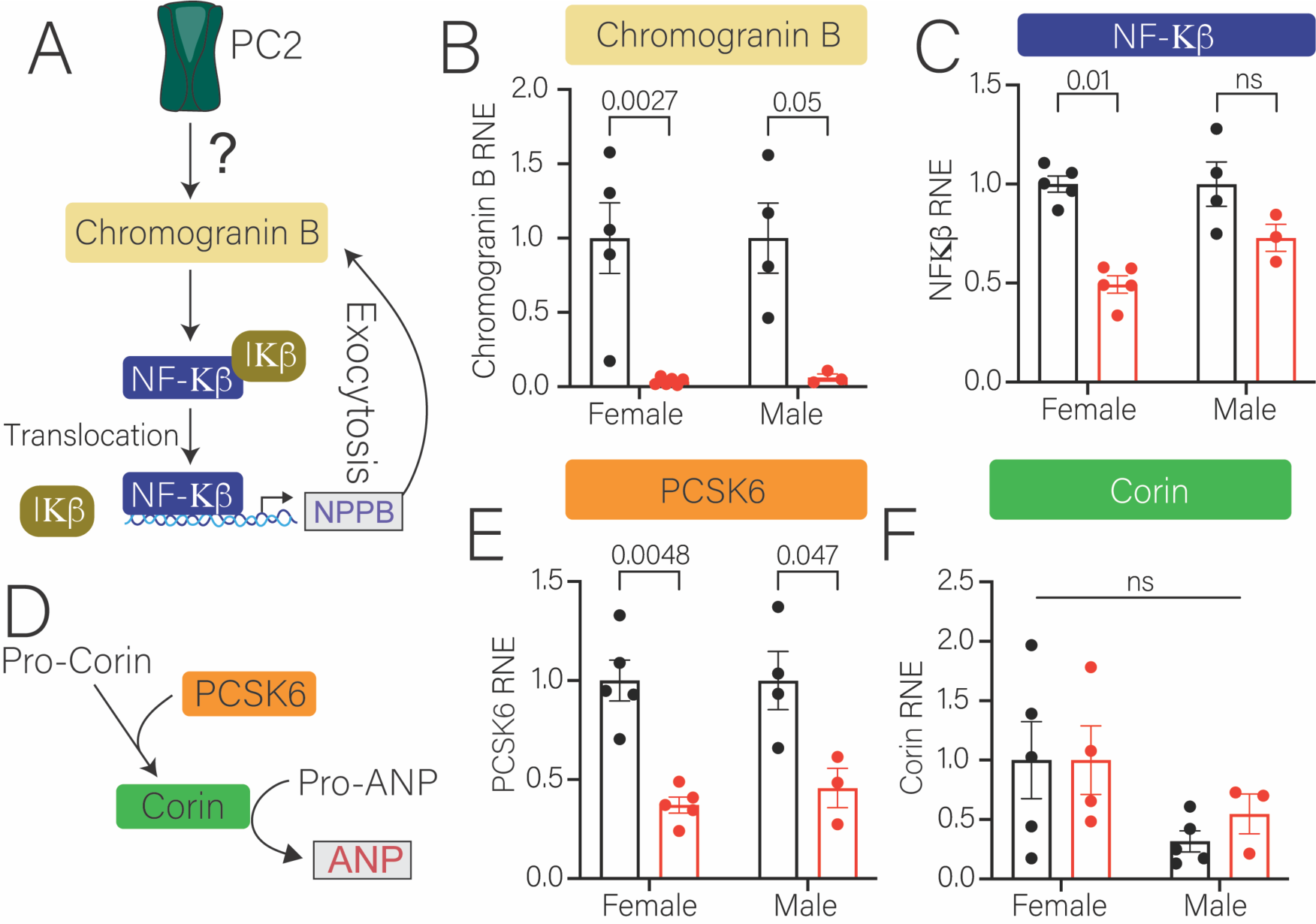
Pathways for NP maturation is decreased in cardiomyocyte specific PC2-KO mice. **A.** Diagram of PC2 regulated BNP production through Chromogranin B pathway**. B.** mRNA expression of Chromogranin B is significantly decreased in tissue from left ventricle in both cPC2-KO female and male mice. Dots represent individual mice. Bars represent mean±SEM. Statistical analysis was done using Kruskal Willis followed by Dunn’s statistical test. p-values are listed in the figure. **C.** mRNA expression of NFκβ is significantly decreased only in tissue from left ventricle of cPC2-KO female mice. Dots represent individual mice. Bars represent mean±SEM. Statistical analysis was done using Kruskal Willis followed by Dunn’s statistical test. p-values are listed in the figure. **D.** Diagram of ANP maturation. **E.** mRNA expression of PCSK6 is significantly decreased in tissue from left ventricle in both cPC2-KO female and male mice. Dots represent individual mice. Bars represent mean±SEM. Statistical analysis was done using Kruskal Willis followed by Dunn’s statistical test. p-values are listed in the figures. **F.** mRNA expression of Corin from left ventricle tissue was unchanged in both female and male control and cPC2-KO mice. Dots represent individual mice. Bars represent mean±SEM. Statistical analysis was done using Kruskal Willis followed by Dunn’s statistical test.

### Constituents of NPR signaling pathways are impaired in cPC2KO mice

qPCR analysis of left ventricular NPR expression showed decreased NPR1 expression in cPC2-KO female mice compared to WT (Supplemental Fig. 4B). Neither mRNA expression of NPR1 in males nor NPR2 expression in either sex were different between WT and cPC2-KO mice (Supplemental Fig. 4C). NPR3 mRNA expression was not present in either WT or cPC2-KO mouse left ventricular tissue (not shown). mRNA expression of NFAT isoforms in LV tissue was analyzed. NFATC2 mRNA was significantly decreased in female cPC2-KO compared to WT mice (Supplemental Fig. 4D), and a decrease in NFATC4 mRNA expression was present in female and male cPC2-KO mice compared to WT (Supplemental. Fig. 4E). Western blot analysis of LV tissue demonstrated decreased phosphorylation of CREB relative to unphosphorylated CREB in cPC2-KO mice compared to WT mice (Supplemental. Fig. 4F). Collectively, these results suggest that despite the presence of hypertension in the cPC2KO, there is little cardiac remodeling as assessed by echocardiography (Supplemental Fig. 1) and no activation of the hypertrophic gene transcription pathway (Supplemental Fig. 4).

**FIGURE 4.**
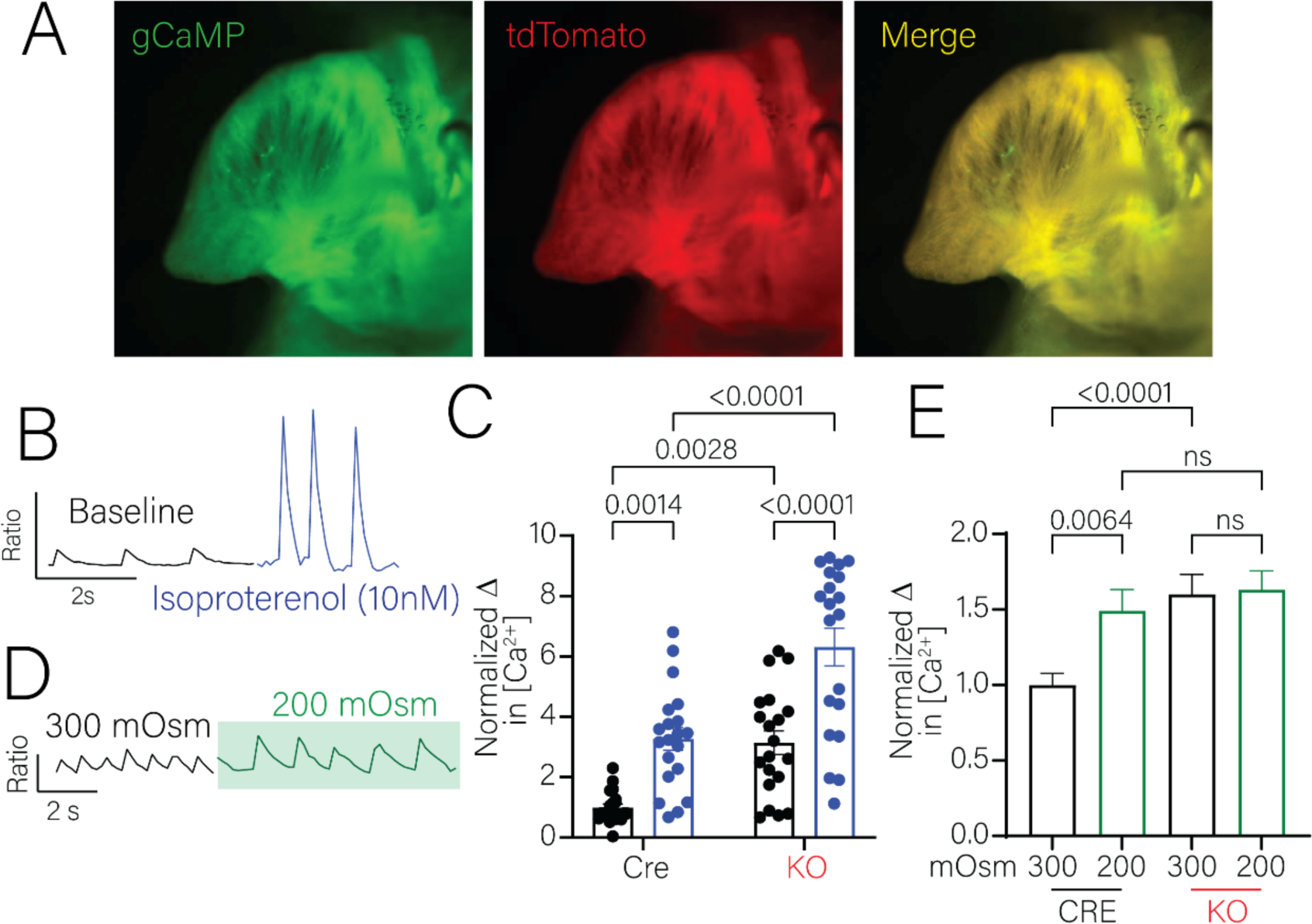
Loss of PC2 results in a deficiency in calcium response to hypo-osmolar challenge in an *ex-vivo* heart model. **A.** Representative images of the right atria of Salsa6f expressing mice showing the expression of gCaMP6F (*left panel*), tdTomato (*middle panel*), and merged image (*right panel*). **B.** Example trace of ratioed spontaneous baseline calcium transients from a control Salsa6F expressing mouse and in response to isoproterenol (10nM, *blue trace*) **C.** Quantified analysis of the normalized area under the curve shows that the cPC2-KO hearts have increased baseline calcium responses compared to control counterparts. In response to isoproterenol, both control and cPC2-KO hearts show statistically significant increases in calcium response. **D**. Example trace demonstrating spontaneous calcium transients at normal cellular osmolality (300mOsm, *black trace*) and in response to hypo-osmotic stress solution (200mOsm, *green trace*). **E.** Quantified analysis of the normalized area under the curve cPC2-KO hearts show increased baseline intracellular calcium compared to control counterparts. In response to 200mOsm hypo-osmotic solution, control hearts demonstrate statistically significant increased intracellular calcium response. cPC2-KO hearts show no significant difference between baseline intracellular calcium and response following hypo-osmotic stimulation. n=4 for control mice and 5 for cPC2-KO mice. Each dot in Panel C represents a region of interest; 5 separate regions per animal were sampled. Bars represent mean±SEM. Statistical analysis was done using 2-way ANOVA. P-values are listed in the figure.

### Impaired Ca^2+^ response in cPC2Kos to hypo-osmotic stimuli

Using a genetically encoded Ca^2+^ sensor mouse expressing td-Tomato and gCaMP6F (Fig. 4A) the cytoplasmic Ca^2+^ response of right atria from WT and cPC2-KO mice were measured during exposure to isoproterenol, (Fig. 4B, C) and decreased extracellular tonicity (200mOsm) (Fig. 4D, E). Before agonist treatment, the area under the curve for cytoplasmic Ca^2+^ was greater in cPC2-KO hearts. Upon addition of isoproterenol, both WT and cPC2-KO right atria increased cytoplasmic Ca^2+^ response similarly (Fig. 4C). When the hearts were exposed to solution at hypo-osmotic conditions of 200 mOsm, the WT right atria demonstrated an increase in cytoplasmic Ca^2+^ which was absent from the cPC2-KO right atria (Fig. 4E). These data suggest that the Ca^2+^ increase associated with hypo-osmotic stretch is separate to the pathways associated with isoproterenol, and that PC2 is in the pathway that contributes to the Ca^2+^ rise associated with hypo-osmotic stress.

### Human iPSC derived cardiomyocytes with PC2 KO have decreased ANP production

Beating ventricular cardiomyocytes derived from human iPSCs were seeded into an engineered heart construct to ensure a regular cellular orientation. After seeding, cells were transduced with lentivirus containing Cas9 and a no-template control (NTC) guide sequence, or a guide sequence against *PKD2* which we had previously verified [60] to create control NTCs and PC2 KOs (Fig. 5A). After maturation, the constructs were inserted into a force-transducer and length controller apparatus (Myopod) to undergo length-dependent activation (LDA) testing from –5% to +5% stretch in 1% increments (Fig. 5A).

**FIGURE 5.**
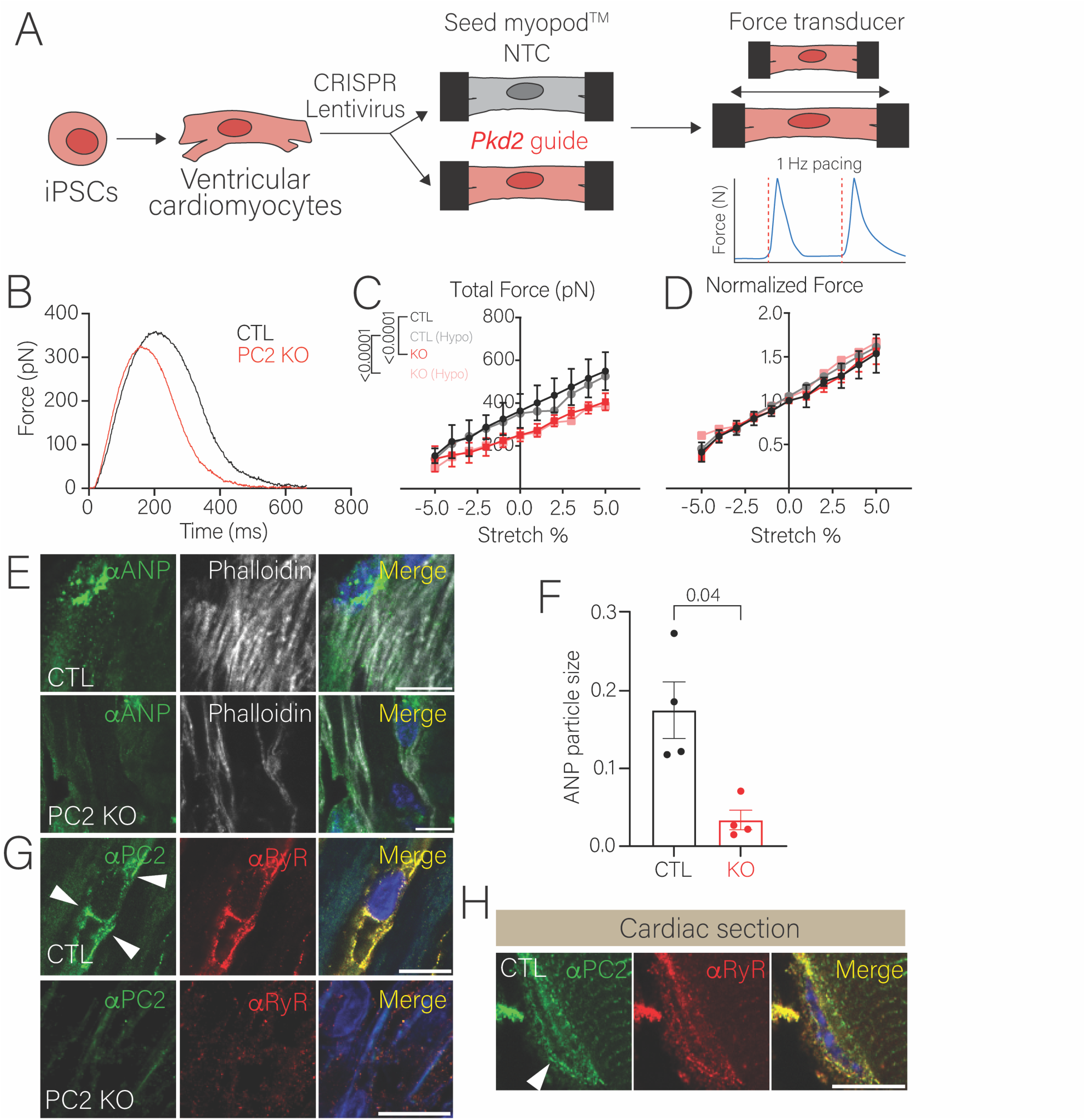
Human iPSC derived cardiomyocytes with *PKD2* KO have decreased ANP. **A.** Schematic diagram of the myopod based experiment. **B.** Example force twitch generated with 1Hz pacing under 0% stretch conditions. **C.** Total force generated with a length dependent activation protocol from -5% stretch to +5% stretch, in CTL (expressing a no-template control guide) and PC2 KO seeded myopods, under conditions of 300mOsm Tyrodes’ and hypo-osmotic Tyrodes’ buffer (270mOsm). **D**. Normalized force curves (to the 0% stretch) demonstrating that neither knockout of PC2 nor hypo-osmotic conditions alter the length dependent activation response. **E.** Example staining of ANP puncta (*green*) in control (*top panels*) and PC2 KO (*lower panel*) following LDA protocol. **F.** Quantification of ANP particle size in CTL and PC2 KO cells demonstrating that the particle size is larger in CTL myopods compared to PC2 KO. Each dot represents example images taken from n=3 myopods per condition. Bars represent mean±SEM. Statistical analysis was done using Mann Whitney U test. p-values are listed in the figure. **G**. Staining of PC2 (*green*) and the ryanodine receptor (*red)* in cells embedded in the myopod. Note decreased PC2 staining in PC2 KO myopods. Scale bar represents 10 µm. **H.** Expression of PC2 around the nuclei in cardiomyocyte sections as well as in close proximity to the ryanodine receptor. Scale bar represents 10 µm.

PC2-KO cardiomyocytes were less able to produce force compared to the NTCs with 0% stretch (Fig. 5B). The total force in relation to stretch at baseline and during hypo-osmotic conditions demonstrated a significant difference between CTL and the PC2-KOs cells (Fig. 5C) which was abolished once the force was normalized to the 0% stretch value (Fig. 5D). This indicates that the loss of PC2 did not impair the degree of LDA of the myofilament and contractility over the range of stretch conditions or with hypo-osmotic stimuli. Immunofluorescence staining for ANP puncta at the completion of the LDA protocol (Fig. 5E) demonstrated a significant reduction in ANP particle size in the PC2-KO cells compared to the CTL cells (Fig. 5F), with similar levels of physical stretch (5%) and hypo-osmotic stimuli. These data are consistent with our animal findings, in that loss of PC2 impairs the production of the NPs. Moreover, loss of PC2 does not impair the ability of the cardiomyocytes to elicit contractile forces suggesting that PC2 functions within a specific sub-domain of a cell separate to the excitation contraction machinery where it facilitates Ca^2+^ release associated with the production of NPs.

To determine if PC2 does reside in a separate domain to the SR associated with the T-tubule, we examined PC2 puncta in both myopod constructs and cardiac sections. We found PC2 localizing in close proximity to the RyR (Fig. 5G, H). However, we also found a separate pool of PC2 surrounding the nucleus, most likely representing the ER.

### Production of NPs is dependent on PC2 but not its calcium activity

Lower mRNA transcription of ANP was demonstrated in PC2-KO cells than in WT cells under both undifferentiated and differentiated conditions (Fig. 6A, B). The re-expression of full-length PC2 in the PC2-KO cell line restored ANP mRNA production (Fig. 6C). The PC2 dead channel, D511V, which lacks ability to conduct ions via PC2 was similarly able to restore ANP production (Fig. 6C). Similarly, the expression of chromogranin B and PCSK6 was restored with either the re-expression of full-length PC2 or the dead channel (Fig. 6D, E). Collectively, these data demonstrates that the presence of PC2 in cardiomyocytes is required for ANP production, but that Ca^2+^ conduction via PC2 is not necessary.

**FIGURE 6.**
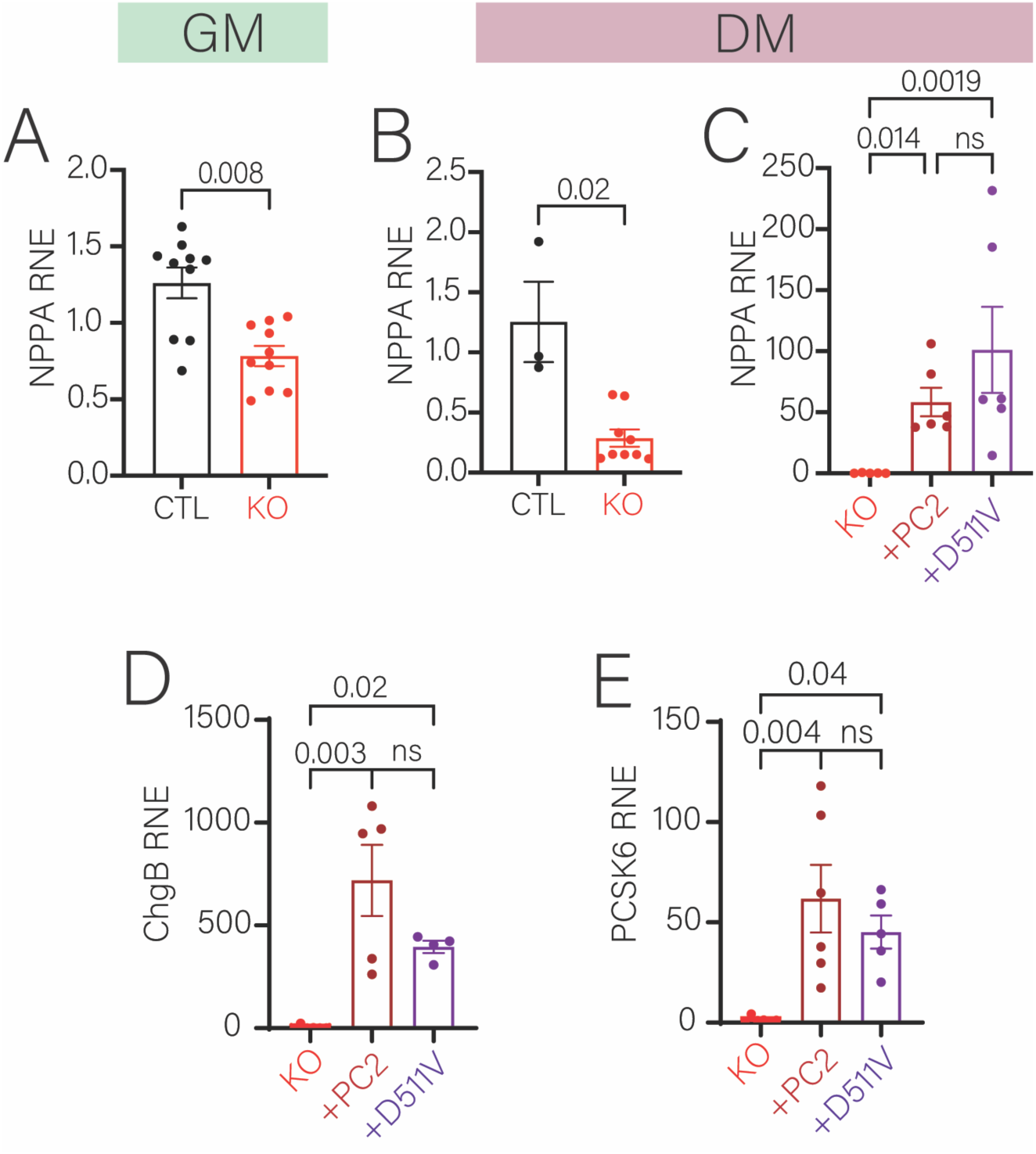
Re-expression of full length PC2 and a non-conducting pathogenic variant restore NPPA expression. **A.** mRNA expression of NPPA is significantly decreased in PC2 KO non-differentiated myoblast C2C12 cells compared to control cells. **B.** mRNA expression of NPPA is significantly decreased in PC2 KO differentiated myoblast C2C12 cells compared to control cells. **C**. Expression of NPPA is significantly increased upon re-expression of either full length PC2 or a pathologic variant which does not conduct calcium. Each dot are biological replicates of the experiment. Bars represent mean±SEM. Statistical analysis was done using Kruskal Willis followed by Dunn’s statistical test. P-values are listed in the figure. **D**. Expression of Chromogranin B is significantly increased upon re-expression of either full length PC2 or a pathologic variant which does not conduct calcium. Each dot are biological replicates of the experiment. Bars represent mean±SEM. Statistical analysis was done using Kruskal Willis followed by Dunn’s statistical test. P-values are listed in the figure. **E**. Expression of PCSK6 is significantly increased upon re-expression of either full length PC2 or a pathologic variant which does not conduct calcium. Each dot are biological replicates of the experiment. Bars represent mean±SEM. Statistical analysis was done using Kruskal Willis followed by Dunn’s statistical test. P-values are listed in the figure.

## DISCUSSION

Over 70% of ADPKD patients have early onset hypertension before decline in kidney function. Here, we demonstrate that PC2 is a key regulator of the cardiomyocyte’s ability to produce NPs and loss of cardiac specific PC2 in the absence of kidney involvement results in elevations in systolic and diastolic blood pressure. We show evidence of impaired NP axis signaling through decreased mRNA expression of ANP and BNP, and a failure to upregulate NP gene expression as evidenced by CgB abolishment and lack of change in GATA4 (Fig. 7). The abolishment of CgB points towards secretory dysfunction of vesicles containing NPs, and decreased PCSK6 indicates an impaired ability to produce functional ANP. Lastly, targets of NPR activity were downregulated, showing an impaired ability to respond to autocrine activity of the NPs. We also demonstrate that human iPSC derived cardiomyocytes lacking PC2 have diminished ANP production. Collectively, we show that there is a multi-pathway dysfunction of the NP system when cardiac PC2 is lost culminating in an impaired ability to respond to increased pressure, sympathetic tone, or tonicity, thus exacerbating a hypertensive phenotype.

**Figure 7.**
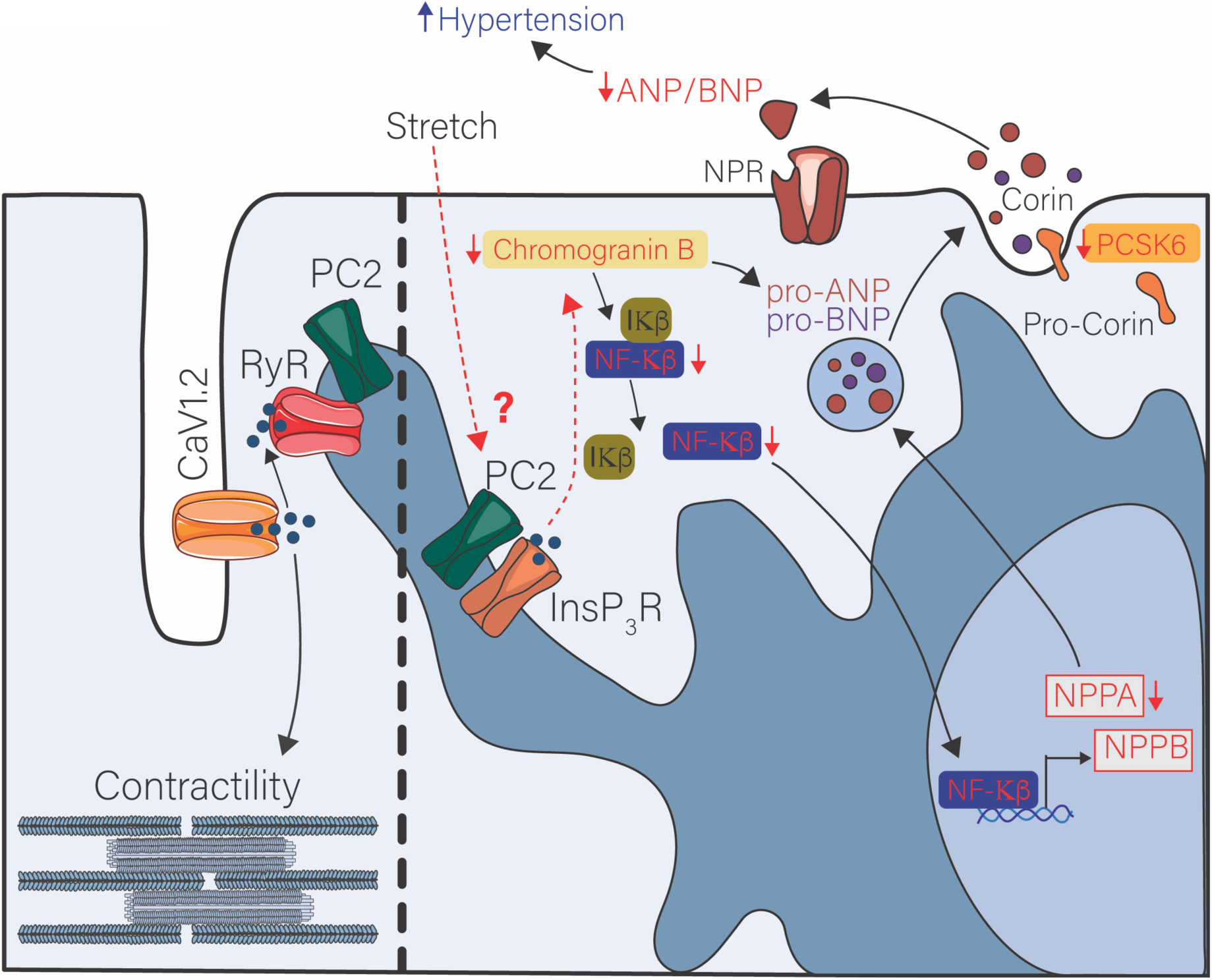
Schematic diagram summarizing the effects of PC2 on the NP pathway. Red letters indicate a decrease while blue letters indicate an increase. Dashed line represents unknown pathway. Small blue circles represent Ca^2+^ ions.

The production of NPs is Ca^2+^ dependent [41,42,44], but the specific molecules responsible for this process have not been identified. *Ex-vivo* preparations of cPC2-KO hearts fail to increase intracellular Ca^2+^ in response to hypo-osmotic stimuli. However, cPC2-KO hearts do exhibit preserved Ca^2+^responses to beta-adrenergic stimuli, indicating that PC2 plays a separate role in osmolality sensing and subsequent signal transduction in the heart. This is consistent with our findings of a dysfunctional NP axis, as hypo-osmolality is a condition that stimulates NP release from cardiomyocytes [22]. As PC2 can conduct Ca^2+^, as well as interact with Ca^2+^ channels including RyR and the InsP_3_R, we used a pore conducting dead mutation of PC2 to test if Ca^2+^ via PC2 was required. As the channel dead mutation, like full-length PC2, was able to restore ANP production, this suggests that PC2 regulates a specific Ca^2+^ pool. Our experiments in human iPSC derived cardiomyocytes suggest that this pool of Ca^2+^ is not linked to excitation-contraction coupling, and thus the most likely candidate is the InsP_3_R surrounding the nucleus [61,62]. This is, however, in contrast to other studies that suggest that Ca^2+^ from the sarcolemma is required for ANP production under hypo-osmotic stimuli [63]. Future studies will examine the exact interactions between nuclear ER Ca^2+^and PC2 in cardiomyocytes.

Our data suggest that ER-PC2 localized away from the sarcoplasmic reticulum contributes to the specific activation of the NP pathway. This contrasts with the localization of the PC1/PC2 complex in renal epithelial cells, where their localization is on the primary cilia, as well as the ER and mitochondrial membranes [64–67]. Unlike renal epithelial and developing cardiac tissue, mature cardiomyocytes apart from the endocardial layer do not have primary cilia [68,69], and thus, a role of polycystins localizing to cardiomyocyte cilia is less likely. Although we only tested the ability of cardiac-PC2 to mediate NP production, cardiac-localized PC1 might have a similar role. Loss of cardiac PC1 from birth results in stunted heart growth and an inability to produce NP with pressure overload [22], suggesting the possibility that PC1/PC2 act in synergistic pathways. Indeed, given that ∼80% of patients with ADPKD have mutations in PC1, and have early onset hypertension, this suggests that PC1 may similarly contribute to cardiac-driven NP pathways like PC2.

Our results suggest an additional extra-renal source of hypertension that may contribute to the early onset hypertension experienced by ADPKD patients. As our cPC2-KO mice did not have renal cysts, it is unlikely that there is impingement of the renal vasculature and activation of the RAAS. Other studies have demonstrated that loss of polycystins in the vascular endothelium results in hypertension [20,70], thus loss of polycystin function in either the heart or endothelium would exacerbate hypertension. However, a deficiency of NP signaling in ADPKD patients may explain their initial hypertensive phenotype. A lack of ability to respond to increased circulating blood volume, blood pressure, or hypo-osmotic stimuli through the same dysfunctional NP pathway could lead to a positive feedback loop, exacerbating and maintaining this hypertensive phenotype (Fig. 6C). Moreover, we observed that the hypertensive phenotype did not itself induce or activate cardiac hypertrophic remodeling genes like NFATC2 or C4. This suggests that PC2 may be a cardiac stress sensor [71], and that the absence of PC2 prevents the engagement of cardiac remodeling.

Hypertension leads to and exacerbates both renal disease and cardiovascular disease, so a lack of resolution of hypertension can contribute to the progression and sequelae of ADPKD. This was shown in the HALT-PKD trial, where patients in the early stages of ADPKD benefitted from rigorous blood pressure control as opposed to standard blood pressure control. Patients in the rigorous control group demonstrated slowing of increases in total kidney volume and declines the left ventricular mass index (LVMI) [12]. LVMI correlates with risk of mortality from cardiovascular disease [72,73].

Current therapies for ADPKD involve the administration of blood pressure control agents, with RAAS inhibitors (ACE inhibitors/Angiotensin Receptor Blockers) being preferred due to their additional nephroprotective effects [74]. Adequate blood pressure control is associated with decreased progression of cardiovascular disease, the main cause of mortality in ADPKD patients. Thus, targeting another mechanism for hypertension, such as an impaired NP response, could provide better control of blood pressure and slow disease progression. Neprilysin inhibitors, such as Sacubitril, are a newer class of medication that inhibit the enzyme responsible for the breakdown of the NPs, Neprilysin. Combined Angiotensin Receptor Blockers -Neprilysin Inhibitors (ARNIs) have had recent FDA approval [75]. ARNIs demonstrate greater therapeutic effects than RAAS inhibitors alone in patients with Heart Failure with Reduced Ejection Fraction (HFrEF), a result of the PARADIGM-HF trial [76,77]. Findings of impaired NP axes in cPC2-KO hearts suggest an alternative explanation for early onset hypertension in ADPKD and warrant further investigation and therapeutic options such as ARNIs. The effectiveness of ARNIs remains to be tested in the ADPKD population. Our study, which demonstrates that human cardiomyocytes lacking PC2 also have diminished capacity to produce NPs, therefore points to a possible primary cardiac etiology for the high incidence of cardiovascular disease, namely hypertension, in models of ADPKD stemming from PC2’s absence.

## CONCLUSION

We show that PC2 plays a multifactorial role in cardiomyocytes’ ability to produce NPs. This is most likely due to PC2’s ability to contribute to Ca^2+^ signals that can amplify downstream signaling to produce NPs. While this study provides another functional role of PC2 plays in the cardiovascular system, it also provides insight into a non-renal source of hypertension that could be seen in patients with ADPKD. As hypertension often arises before loss of renal function occurs in ADPKD patients, finding a non-renal mechanism for this hypertension provides the possibility of early intervention, more targeted therapies, and staving off future complications.

## ACKNOWLEDGEMENTS

Research reported in this publication was supported by the National Institute of Diabetes and Digestive and Kidney Diseases of the National Institutes of Health under Award Numbers U2CDK129917 and TL1DK132769 (KMMN), NHLBI T35 HL120835 (BE); R00DK101585 and PKD Foundation 1021282 (IYK) and R00141698 (DYB). Pilot funds from UL1TR002389 (IYK and ABC) are acknowledged. We acknowledge NIH funds 1S10OD028449 to purchase the VEVO3100. We thank the Department of Cell and Molecular Physiology (Loyola University Chicago) for access to the specialized imaging resource center (Zeiss 880-Airyscan) and the Loyola CVRI support for telemetry equipment and the iPSC resource.

## Contributions

Conception: BL and IYK. Experiments: BL, KMMN, PT, ED, RMK, GF, QC, MW, IYK. Analysis: BL, KMMN, MW, IYK. Discussions: BL, KMMN, RMK, QC, ABC, DYB, IYK. Draft: BE, KMMN and IYK. All authors approved the final manuscript.

## Conflicts of interest

Authors have no conflicts.

## SUPPLEMENTAL FIGURE LEGENDS

**SUPPLEMENTAL FIGURE 1.**
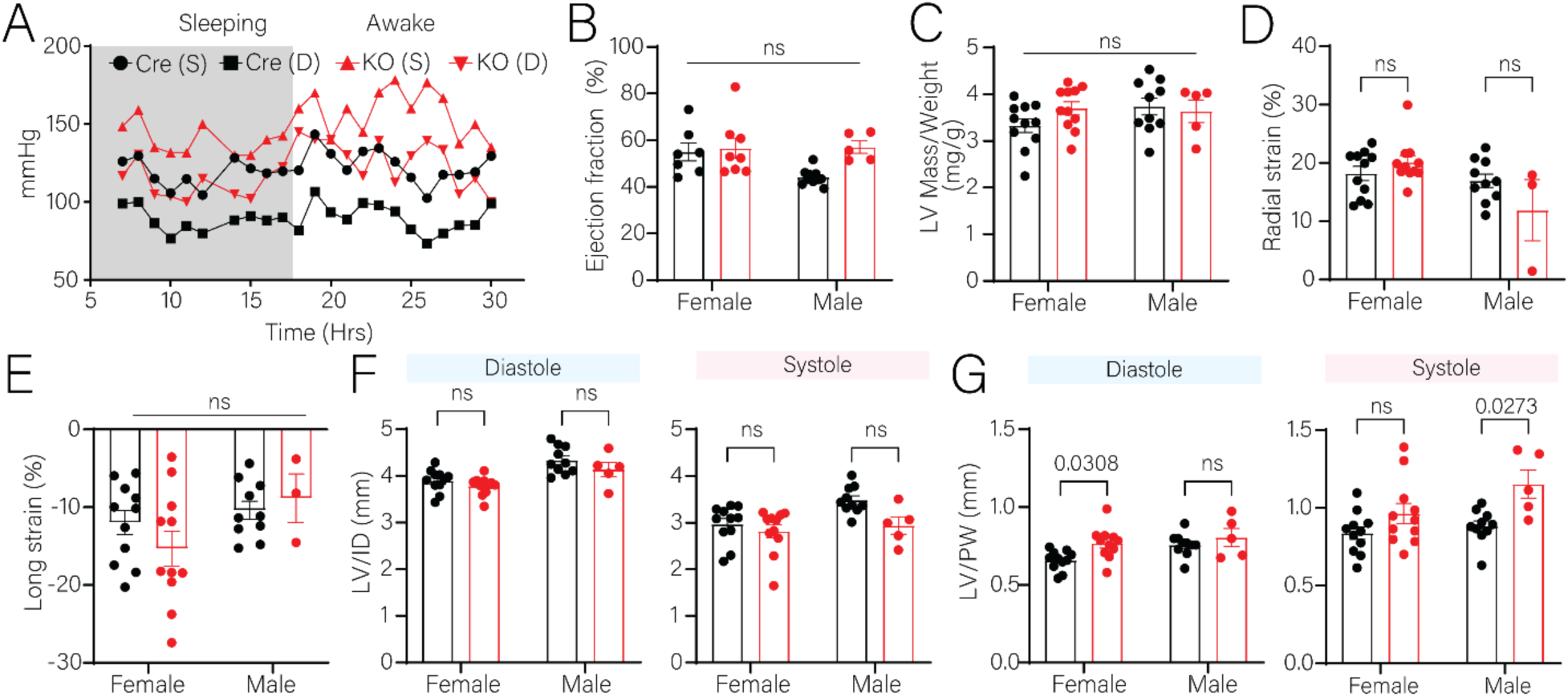
**A.** Representative tracing of blood pressure in male control and cPC2-KO mice for a period of 30 hours. **B.** Ejection fraction (EF) is unchanged in male and female mice. **C.** Left ventricular (LV) mass is unchanged in male and female mice. **D**. Global radial strain is unchanged. **E**. Global longitudinal strain is unchanged. **F.** The inner diameter (ID) of the left ventricle is unchanged under diastole or under systole. **G**. The posterior wall (PW) is thicker in female under diastole but unchanged in males. The posterior wall (PW) is thicker in males under systole but unchanged in females. Dots represent individual mice. Bars represent mean±SEM. Statistical analysis was done using Kruskal Willis followed by Dunn’s statistical test. p-values are listed in the figure.

**SUPPLEMENTAL FIGURE 2.**
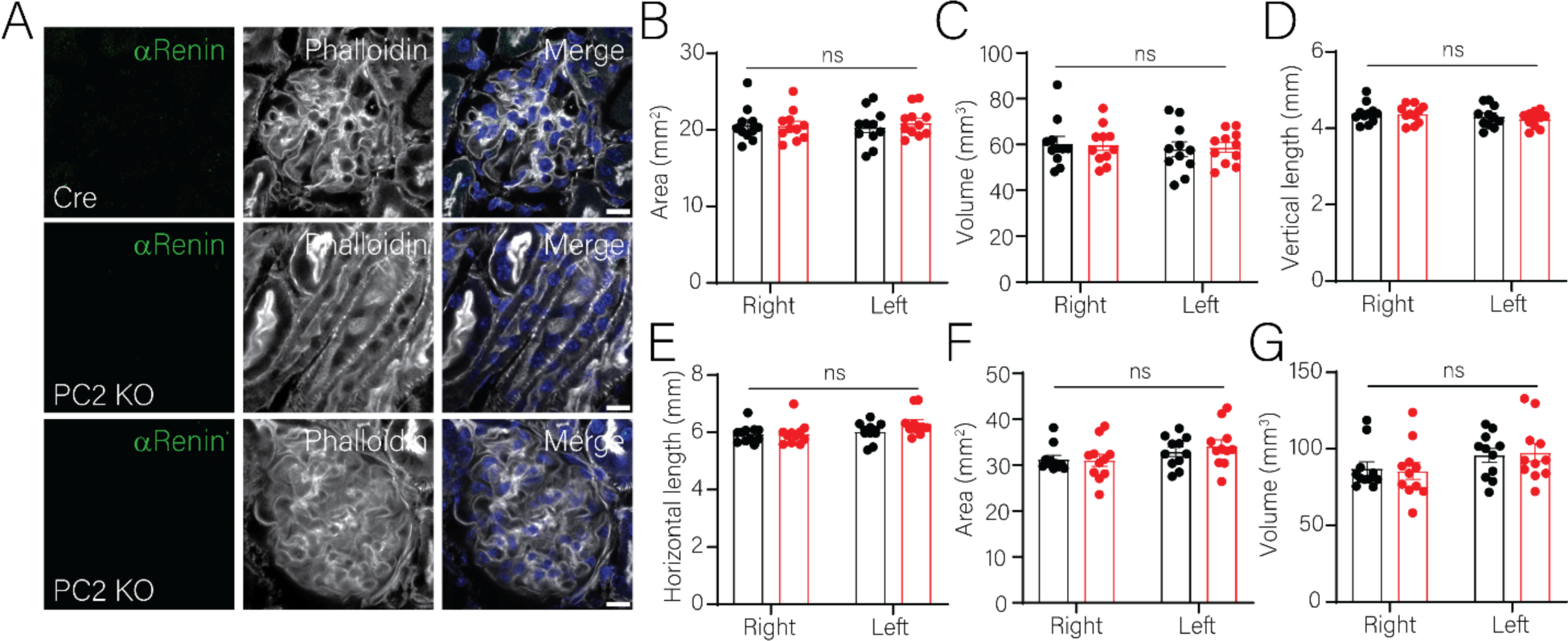
**A**. Representative immunofluorescent images of renin expression (*green*) which is not evident in either CTL or cPC2-KO mice. cPC2-KO mice do not have tubular cysts. **B-G**. All kidney parameters as measured by ultrasound are not significantly different between control and cPC2-KO mice (female mice tested).

**SUPPLEMENTAL FIGURE 3.**
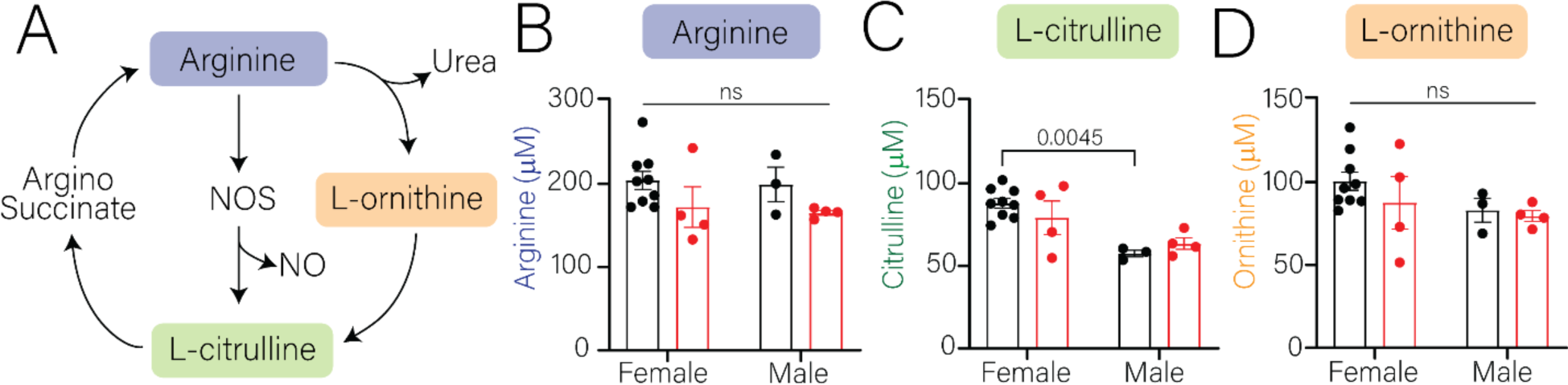
**A**. Diagram of metabolic arginine to citrulline pathway. **B-D**. Arginine, L-citrulline and L-ornithine in the serum of cPC2-KO and CTL mice are not different.

**SUPPLEMENTAL FIGURE 4.**
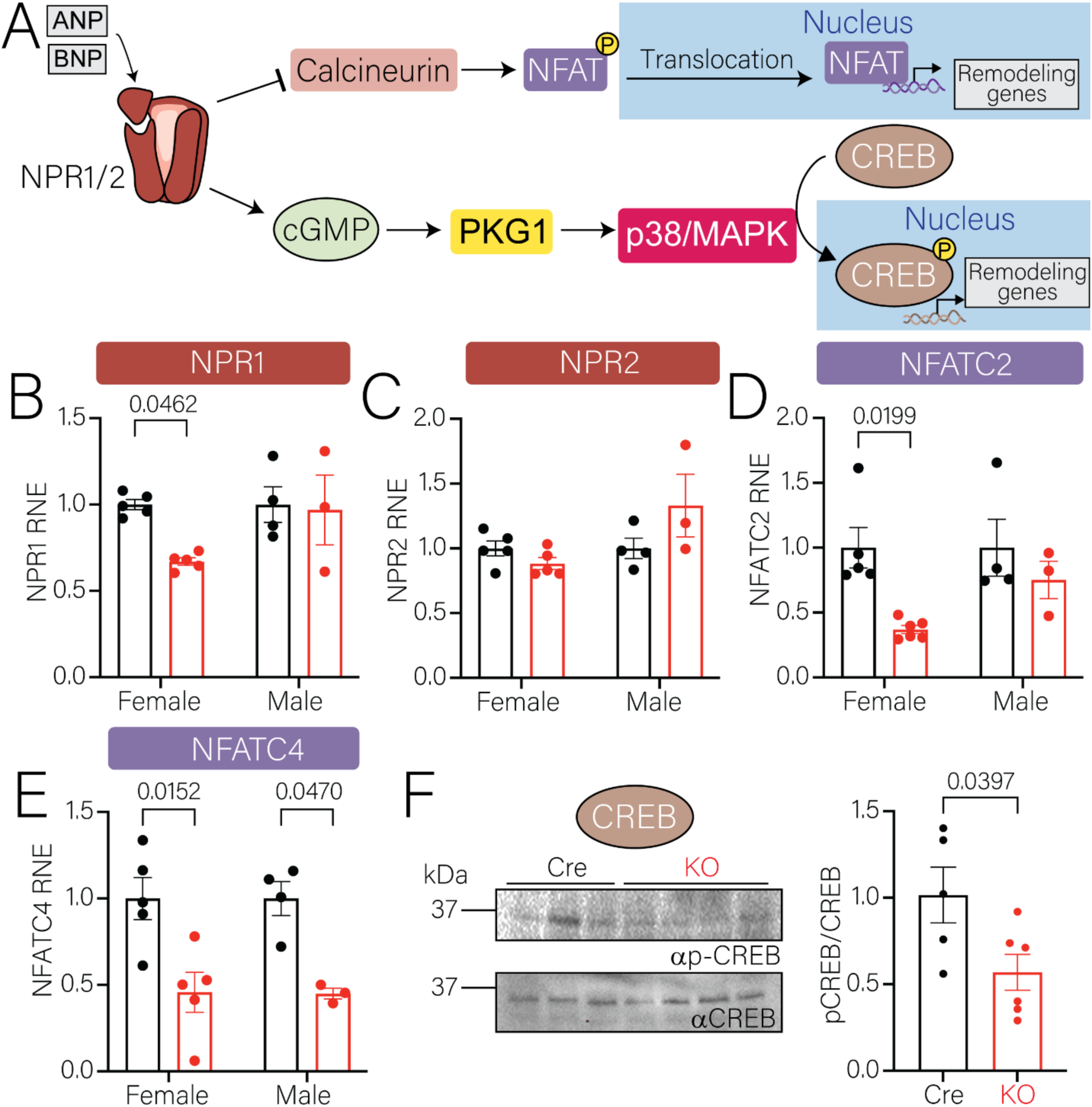
**A.** Schematic diagram of downstream ANP/BNP autocrine signaling pathways. **B.** Decreased NPR1 mRNA expression in female cPC2-KO mice compared to control mice. No difference between males. Each dot is a separate animal. Bars represent mean±SEM. Statistical analysis was done using Kruskal Willis followed by Dunn’s statistical test. P-values are listed in the figure. **C**. No change in NPR2 mRNA expression between cPC2-KO mice and control mice, both females and males. **D**. Decreased NFATC2 mRNA expression in female cPC2-KO mice compared to control mice. No difference between males. Each dot is a separate animal. Bars represent mean±SEM. Statistical analysis was done using Kruskal Willis followed by Dunn’s statistical test. P-values are listed in the figure. **E**. Decreased NFATC4 mRNA expression in female and male cPC2-KO mice compared to control mice. Each dot is a separate animal. Bars represent mean±SEM. Statistical analysis was done using Kruskal Willis followed by Dunn’s statistical test. P-values are listed in the figure. **F**. Western blot of pCREB and CREB from LV tissue in control and cPC2-KO mice. **G**. Quantification of western blots. Each dot is a separate animal. Bars represent mean±SEM. Statistical analysis was done using Mann Whitney U test. p-values are listed in the figure.

**SUPPLEMENTAL DATA on request**: Metabolomic data

## Notes

### Competing Interest Statement

The authors have declared no competing interest.

